# Spatially explicit sampling frameworks to identify regions of increased mosquito abundance

**DOI:** 10.1101/2023.11.21.568008

**Authors:** Brigid Kemei, Eric Ochomo, Maurice Ombok, Janet Midega, Eric R. Lucas, Martin J Donnelly, Luigi Sedda, Daniel P. McDermott

**Author notes:** **Contact information:** Brigid Kemei, Eric Ochomo, Maurice Ombok;, Janet Midega, Eric R Lucas, Martin J. Donnelly, Luigi Sedda, Daniel P. McDermott. **Corresponding Authors:** Eric Ochomo, Brigid Kemei, Daniel P. McDermott, **Full address**: Department of Vector Biology, Liverpool School of Tropical Medicine, Pembroke St, Liverpool L3 5QA.

## Abstract

Vector control interventions often lack comprehensive information on vector population distribution and dynamics. This knowledge gap poses challenges in targeting interventions effectively, especially in areas with heterogeneous transmission and where complementary vector control tools may be required to achieve sustained impact on disease transmission. In this study, we implemented a spatially explicit sampling framework for improved vector surveillance in coastal Kenya. Our stratified lattice with close pair sampling design aimed to characterise the vector dynamics of the primary malaria-transmitting species in the area and assess the ecotype classification’s effectiveness at identifying clear population patterns. The study collected 3,621 mosquitoes, with *An. funestus* s.l. being the most abundant malaria vector. The inclusion of the ecotype classification significantly improved spatial abundance model estimates for *An. gambiae* and *Culex* spp. Wetlands, topographic wetness index, and proximity to rivers were associated with increased mosquito abundance. Spatial modelling revealed high abundance regions near the Galana-Sabaki River. Our study demonstrates the applicability of a reproducible spatial sampling approach to identify areas with high vector abundance and inform targeted vector control strategies. The study highlights the importance of ecological stratification and a spatial explicit sampling approach for predicting mosquito presence when prior data is limited and underscores the potential for refining future sampling for control efforts.

## Introduction

The decision to deploy a specific vector control intervention at a sub-national level is often made with limited information on the distribution and dynamics of the vector population in the intervention area (Ozodiegwu et al. 2021; Walker et al. 2016; Burkot et al. 2019). This reflects the challenges around maintaining surveillance systems that identify regions of high transmission potential and the presence of resistance mechanisms that may impede control measures (Russell et al. 2020). A highly granular understanding of vector distribution and bionomics may not be seen as a pressing requirement if interventions can be scaled, at limited additional cost, to large areas such as the deployment of LLINs. However, there is growing recognition that as prevalence decreases and transmission becomes more spatially heterogeneous, complementary, focal-tools are needed to sustain impact in a cost-effective manner (Barreaux et al. 2017; Killeen et al. 2017; Killeen 2014). These supplementary tools often require targeting to make them financially/logistically viable (e.g. larviciding) and they, therefore, require a much greater degree of surveillance data (Chaki et al. 2009). In areas with limited prior data, there are no clear guidelines on establishing a powered prospective sampling programme that can both guide vector control operations and inform a suitable surveillance strategy capable of providing key evaluation metrics.

Entomological research often relies on selecting sampling sites based on the specific goals of the study. However, a common obstacle in implementing regular monitoring lies in determining the locations for sampling. They could conduct simple random sampling to ensure unbiased independence between sampling sites, but this method may not provide a representative sample of the area given, for example, underlying differences due to ecotype variation or clustering of points (Inman et al. 2021; McGarvey, Burch, and Matthews 2016)). If a study requires large numbers of mosquitoes for insecticide resistance monitoring, researchers may sample in locations that have provided large numbers in the past (legacy sampling). Similarly, collection site accessibility can be a factor in sampling location selection (convenience sampling) (Longbottom et al. 2020). Therefore, collating data sets from several projects coupled with limited information on the sampling methodology may lead to a biased interpretation of vector dynamics in the area (Massey et al. 2016; Zhou et al. 2009; Service, M. W. 1977). There have been calls for more rigorous surveillance frameworks, that capture the range of eco-epidemiological settings, for the routine sampling of mosquito populations. The recent development of the Entomological Surveillance Planning Tool provides clearer guidance on the key information required to establish a monitoring network (World Health Organization 2018; University of California, San Francisco. 2022).

The sampling approach deployed by the National Malaria Control Programme in Kenya (NMCP) in Kenya uses malaria data collated by health facilities to identify three sub-counties within each County that are classified as having, in relative terms, high, medium and low transmission. Entomology surveillance teams then conduct convenience sampling within the selected sub-counties to determine the vector species present and their insecticide susceptibility profiles (National Malaria Control Programme 2019). There is a recognition of the need to improve the sampling frameworks both to capture emergent threats such as the recent discovery of invasive *Anopheles stephensi,* but also to understand how vector bionomics vary over space and time (Ochomo et al. 2023; Ndenga et al. 2023; Mnzava, Monroe, and Okumu 2022).

In this study, we trial the application of a spatial explicit sampling framework for improved vector surveillance in coastal Kenya (Sedda et al. 2019). The coastal region of Kenya has seen a period of a marked decrease in malaria morbidity between the years 1998 and 2011. Recent studies in Kilifi in 2019 found that the all-age *Plasmodium falciparum* prevalence was ≈∼10% in contrast to the >40% reported in 1998 (O’Meara et al. 2008; Snow et al. 2015; Kamau et al. 2020; Mogeni et al. 2016).

In this study we have deployed a stratified lattice with a close pairs sampling design to identify, with associated uncertainty, primary malaria vector abundance across the study area in coastal Kenya, to assess the strength of the ecotype classification in capturing differences in vector abundance and to characterise the presence of secondary *Anopheles* spp. in the region.

## Methods

### Study area

This study was conducted near Malindi town in the north of Kilifi County, a malaria-endemic coastal region of Kenya. The area has a tropical dry savannah climate with mean maximum daily temperatures ranging from 27-32°C. There is perennial malaria transmission in the region and a number of important *Anopheles* species have been previously identified including *Anopheles gambiae s.s*, *An. arabiensis*, *An. merus* and *An. funestus* (Mbogo et al., 2003). Resistance to pyrethroids, carbamates and organophosphates has been reported in the region (Ondeto et al. 2017; Munywoki et al. 2021). Kilifi County receives the LLINs as part of the national mass distribution campaign with high reported usage at all ages (>70%) (Kamau et al. 2022).

### Sampling

The sampling area is described in detail in (Sedda et al. 2019) and shown in Figure 1. In brief, the area was stratified into ecotypes using a supervised spatial quadratic discriminant analysis which utilised a number of widely available data sources such as land cover (GlobeLand30), climate (MODIS Enhanced Vegetation Index; MODIS Air Temperature; MODIS Evapotranspiration; WorldClim Version 2 Precipitation) and elevation (NASA Shuttle Radar Topographic Mission (SRTM) 90m) variables. The optimal number of ecotypes is selected based on the Wilks’ criterion (el Ouardighi, el Akadi, and Aboutajdine 2007; Sedda 2023).

**Figure 1.**
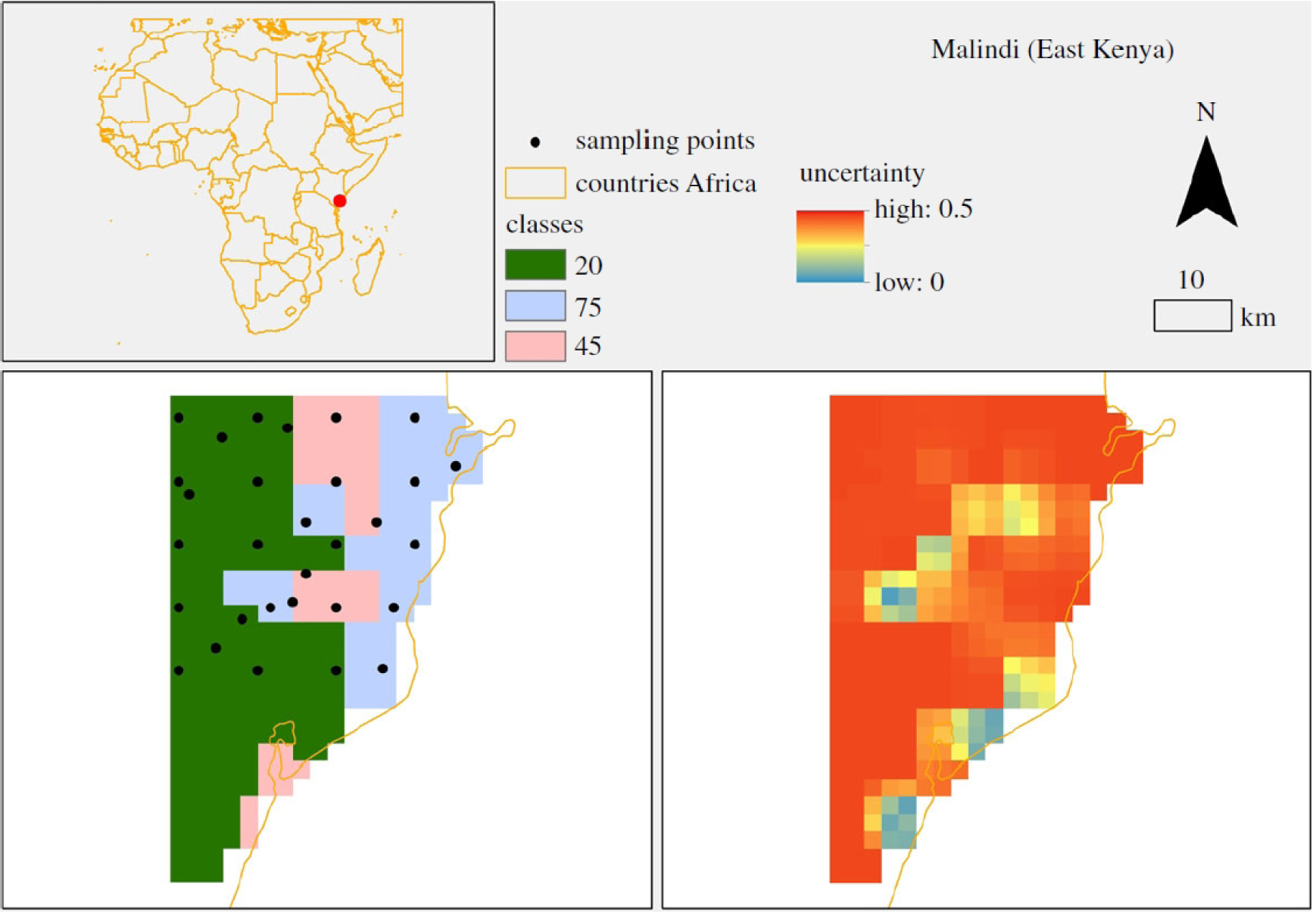
Sampling framework developed by Sedda *et al*. (2019) and implemented in the present study (Reproduced with permission). Lower maps are ecological classification (left) and the associated uncertainty in classification (right) for the area of Malindi. Classes are 20= forest; 75=urban; 45=scrubland. Map was made using ArcMap 10.4 (http://desktop.arcgis.com/en/arcmap/). Source administrative limits: http://www.maplibrary.org/library/index.htm. The true sampling locations differ, as outlined in Figure 2, reflecting where the nearest group of structures were identified from these predefined sampling points.

While the classification classes are identified primarily according to their dominant land cover class, they are a mixture of all of the above (cover, climate, topography). These ecotypes encompass the variation across all these datasets but for ease, they will be referred to by the dominant land use within each category: forest, scrubland and urban (Figure 1; Supplemental Table 1). A representative stratified lattice with a close pairs sampling design was generated with the number of sampling locations in each ecotype weighted by the proportion of the area covered by each ecotype. Close pairs were allocated at random with the condition that they are at least 1 km distant from the sampling locations in the lattice. Collections were carried out at thirty locations (sample size derived from (Sedda et al. 2019)) every fortnight for two months between 1^st^ October and 18^th^ November 2018. Upon arrival at the GPS-guided sampling points, the team identified the nearest group of houses within a 5km radius (Figure 2). Four households at each site were selected at random and consent was obtained. One of the four household heads was selected and trained to assemble and operate a CDC light trap. Light traps were placed indoors near to sleeping space and were run overnight for approximately 12 hours. All mosquito samples were stored in individual Eppendorf tubes and preserved using silica gel for morphological species identification and sexing at KEMRI-Center for Global Health Research, Kisian.

**Figure 2.**
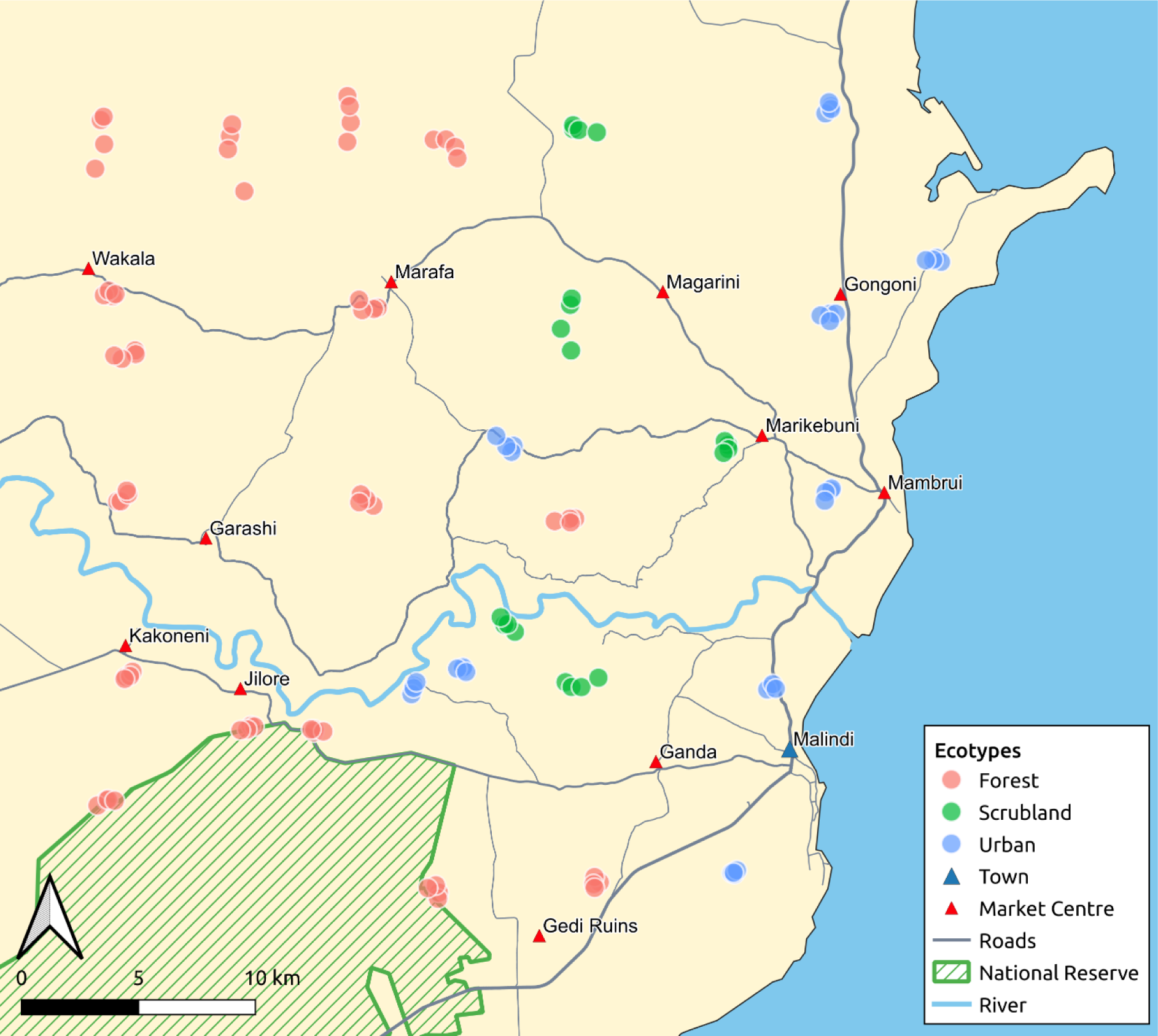
The sampling locations for the lattice with close pairs design implemented in Malindi, Kenya. The sites are arranged in a grid with close pair points added to be representative of short-scale effects (shorter than the lattice resolution). Each ecotype (Forest, Scrubland, Urban) contained a number of sampling locations proportional to its size. When a sampling location fell in an uninhabited area, the sampling team selected the nearest households available.

### Covariates

The framework was designed to ensure sampling effort was proportionate to the area covered by the three ecotypes. To allow for the fine-scale prediction of abundance four additional variables that can be associated with the bionomics of the mosquito populations were screened for predictive ability of the following:

1. We calculated the number of buildings present within a 1km radius of the sampling point as a proxy for both the population of potential hosts and the extent of the built environment. These data were derived from satellite imagery at 100m (Dooley et al. 2020).
2. The topographic wetness index estimates soil wetness using a combination of Shuttle Radar Topography Mission elevation and slope in conjunction with catchment areas to identify points with an increased likelihood of water accumulation (Hengl et al. 2015; Vagen 2010). This has been previously identified as a significant predictor of malaria infection in the western Kenyan highlands (Cohen et al. 2010, 2008).
3. The straight-line distance between the sampling location and river network with the assumption that it may provide riparian breeding sites for mosquito larvae (OCHA ROSEA 2003).
4. While land cover was included in the generation of the ecotypes, land use types that were not included in the initial categorisation were extracted from the annual Copernicus global land change maps for 2018 with the presence or absence of herbaceous wetland or land involved in agricultural production within a 1km radius of the collection point (Buchhorn et al. 2020).

### Statistical analysis

Our primary intent was to highlight regions with higher mean abundances of each species and to determine whether the ecotype categorisation we generated was significantly related to the abundance. We used a generalised linear mixed model (GLMM) with a negative binomial error distribution (to account for overdispersion in the count data) to test the relationship between the number of mosquitoes captured in the households and the covariates outlined above (Harrison et al. 2018). A random effect was included at the household level to account for the repeated sampling in the same household across the period. Models were run for the two primary malaria vectors in the area, *An. gambiae s.l.* and *An. funestus*; and for *Culex* spp which has been implicated in its role as a nuisance biter in intervention usage patterns (Irish et al. 2008; Hill, Lines, and Rowland 2006). The variance inflation factor was assessed for each model with parameters having a VIF greater than 3 being dropped from the model to remove potential collinearity in predictors (Zuur, Ieno, and Elphick 2010). The significance of the ecotype variable was assessed using a log-likelihood ratio test between a full model containing and not containing it using the lmtest package in R (Zeileis and Hothorn 2002). Post-hoc multiple pairwise comparisons of the ecotype levels were carried out using Tukey’s all-pair comparison using the multcomp package with a Holm-Bonferroni correction to account for multiple testing (Hothorn, Bretz, and Westfall 2008). Whilst the sampling framework can be used to capture heterogeneity in space and time, this first deployment of the sampling framework was conducted over a relatively small temporal window so we aggregated the abundance data across the 4-time points (2 months of sampling) to give the total collected in each household over the entire sampling period. We assessed for spatial autocorrelation in the residuals of the models by Moran’s I using the DHARMA package (Hartig 2021). We generated predictive abundance maps for the three species groups we modelled using the sdmTMB package which allows for the incorporation of the functionality of the r-INLA package to approximate a Gaussian Markov random field (GMRF) using a stochastic partial differential equation (SPDE) mesh to account for spatial autocorrelation within a GLMM framework (Anderson et al. 2022; Dormann et al. 2007; Lindgren and Rue 2015). Owing to the challenge that spatial autocorrelation can pose on parameter estimation, significant variables from the non-spatial GLMM were noted and carried forward to generate 3 model sets (a spatial intercept-only model, a significant variable-only model and a maximal model which contained all the initial covariates including those found to be non-significant) (Rousset and Ferdy 2014; Mets, Armenteras, and Dávalos 2017). In the absence of subsequent datasets to conduct validation, k-fold cross-validation (k=10) was carried out on each model and compared by predictive log-likelihood for each species (Yates et al. 2023). Total abundance prediction estimates for the optimal model were generated at a resolution of 1km^2^ together with an associated standard error.

## Results

A total of 3,621 mosquitoes were collected with 72.08% (n=2610) identified as *Culex* spp. with the remainder *Anopheles* spp. The most abundant *Anopheles* was *An. funestus s.l.* (n= 689; 68.11%) found at 16 of the 30 sites, other *Anopheles* included *An. gambiae s.l.* (n = 137; 13.52%) at 17 sites, *An. coustani* (n = 97; 9.58%) at 6 sites, *An. pretoriensis* (n=49; 4.84%) at 5 sites and *An. pharoensis* (n= 33; 3.26%) at 3 sites. *An. squamosus, An. moucheti,* and *An. maculipalpis* were also collected but constituted less than 1% of collected samples. A single sample site on the border of the Arabuko Sokoke National Reserve recorded the presence of all eight *Anopheles* spp.This contrasts with the *Culex* population which, while displaying heterogeneity in abundance, was ubiquitous across the sampling area.

(Figure 3). For subsequent analyses, we focus on the two primary malaria vectors and the abundant *Culex* spp. The overall mean nightly collection per house of *An. funestus* was 1.44±0.3(SE) compared to 0.286±0.05(SE) in *An. gambiae* and 5.44±0.4(SE) in *Culex* spp, with a large degree of variability between the ecotypes (Figure 4).

**Figure 3.**
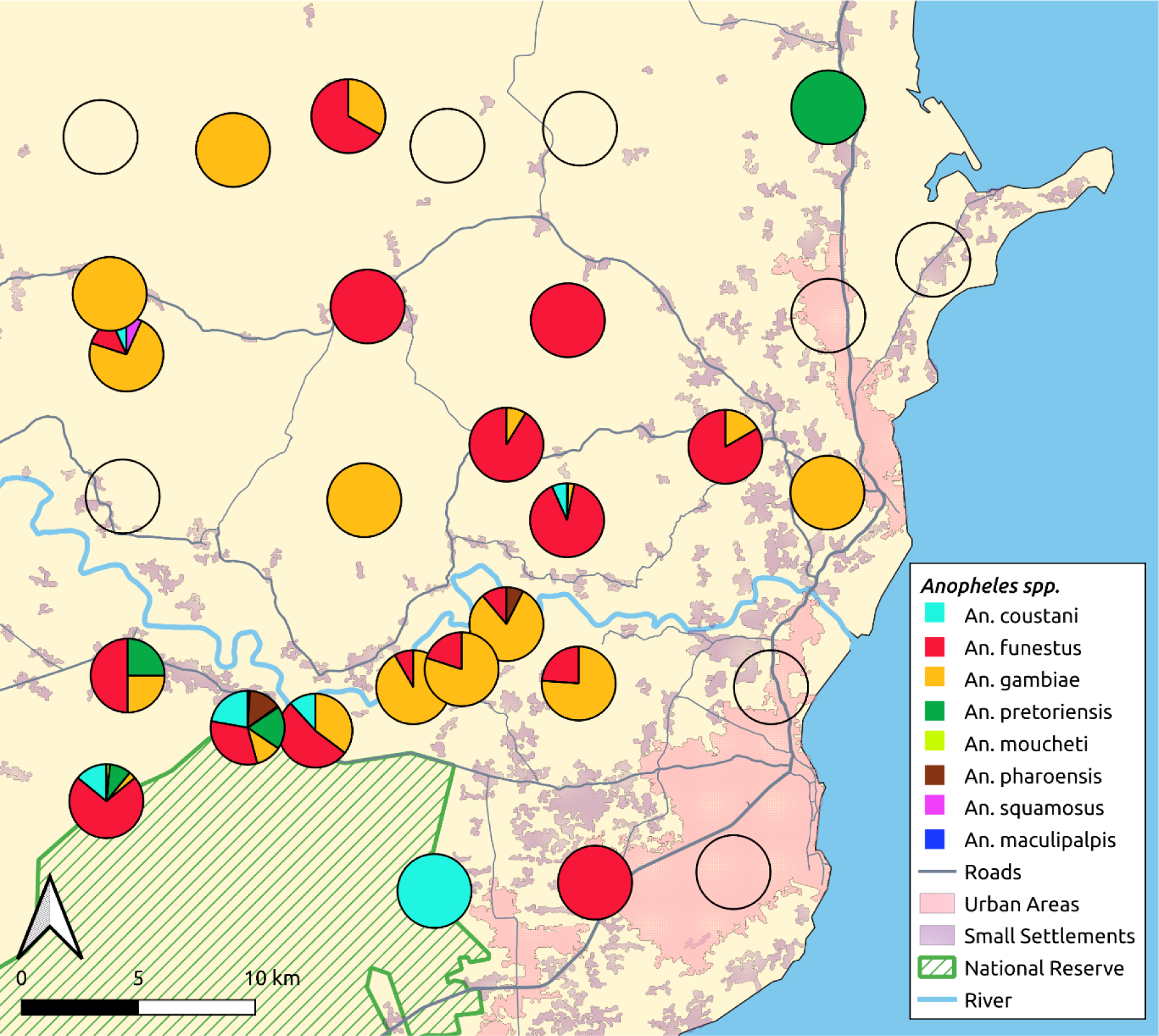
Relative abundance of *Anopheles* species at each of the 30 sampling points in coastal Kenya aggregated across the study period. Empty circles indicate no *Anopheles* spp. mosquitoes were captured in that sampling point across the time period. The region in hatched green indicate the Arabuko Sokoke National Reserve. Urban built up areas are defined as an areas greater than 0.4km^2^ with an overall density >13 households. Small settlements are building groups <50 that do not meet the built up area definition.

**Figure 4.**
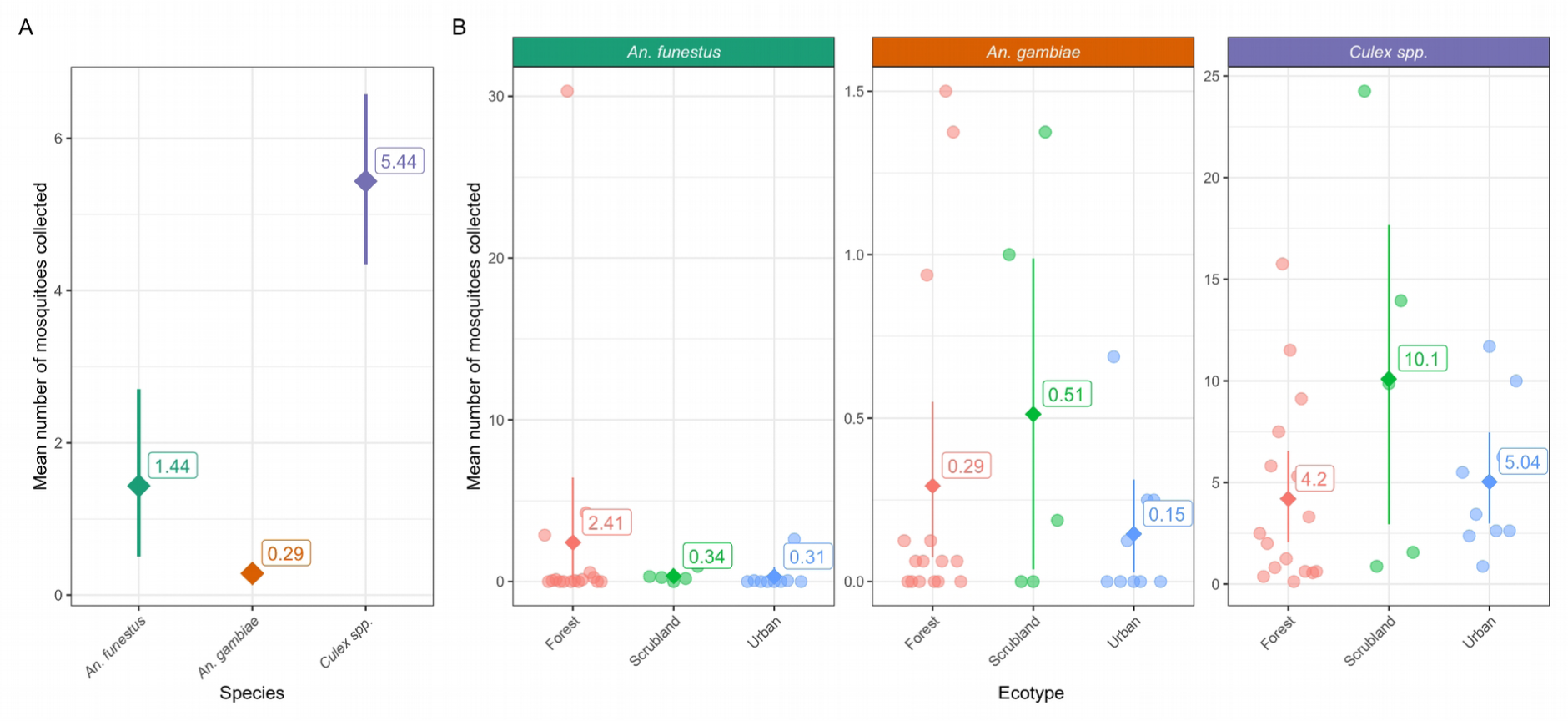
Mean collection per trapping night of *An. gambiae*, *An. funestus* and *Culex* spp. with 95% confidence intervals. (A) The mean number of each species captured each trapping night across the entire sampling area. (B) The mean collection of the 3 species group for each sampling location (●) and each ecotype per trapping night (◆).

When modelled to account for the repeated sampling of the households across the study period and the addition of the environmental covariates, the inclusion of the ecotype did not lead to significant improvement in the model for *An. funestus* (χ2(2) = 3.2745; p =0.1945) but did for *An. gambiae* (χ2(2) = 7.5204; p = 0.02328) and *Culex* spp. (χ2(2) = 10.152; p = 0.006246). Subsequent post-hoc comparison of the ecotypes within *An. gambiae* model found that the Scrubland (x̄ = 2.41; Z = 2.773; p = 0.0151) had a significantly higher mean catch than the Urban ecotype (x̄ = 0.3). Similarly, *Culex* spp. were most commonly found in the scrubland ecotype (x̄ = 10.1) which was significantly greater than the mean collections in the Forest (x̄ = 4.2; Z = 2.9; p =0.01) and Urban ecotypes (x̄ = 5.04; Z = 2.866; p = 0.01). The explanatory variables are derived from open datasets that describe the environmental conditions immediately surrounding the sampling location at a spatial resolution of 1km or less. As a result, there was limited capacity to account for household variation which was substantial across the study area with an estimate of the standard deviation of the random effect variance for the household ranging from 2.966 in *An. funestus* to 1.011 in *Culex* spp. and 0.8226 in *An. gambiae*. The only variable that was significantly associated with the increased abundance of all three species was the presence of wetlands within 1km of the sampled household (Figure 5). In *An. gambiae* there is also a significant positive association with areas that have greater potential for water accumulations (TWI - IRR = 1.35; p = 0.004) which could act as breeding sites. Mean housing density was the only other significantly associated variable with the *Culex* spp. (Mean housing density - IRR = 1.25; p = 0.005). The maximal model residuals for all three species groups, *An gambiae* (Moran’s I = 0.14; p = <0.001), *An funestus* (Moran’s I = 0.19; p =<0.001), and *Culex* spp. (Moran’s I = 0.295; p = <0.001), were found to contain significant spatial autocorrelation. As such we utilised spatial modelling to account for this. The spatial range of the maximal models derived from the Gaussian Markov random field, which in this context is defined as the distance when the spatial correlation between sampled points decays to below 0.13 and denotes the distance in which we may consider spatial autocorrelation to have sufficiently diminished to be deemed as absent (Bakka et al. 2018; Lindgren and Rue 2015), was largest for *An. gambiae* 13.16km (95%CI 4.56 - 37.96km) compared to 11.4km (95%CI 5.29 - 24.6) and 7.36 (95%CI 3.98-13.6) for *An. funestus* and *Culex* spp. respectively (Supplementary Table 2). From the spatial predictions, all three species share a region of high abundance in the centre of the sampling area which co-locates with an area of wetland and proximity to the river course (Figure 6). The comparison of the predictive capability of the model was assessed by k-fold cross-validation to identify the optimal model for generating the abundance surface plots using the lowest value of the negative log-likelihood as an estimate of predictive ability in out-of-sample test folds. For *An. gambiae,* the maximal model was best whereas for *An. funestus* the significant-only model produced the best predictive estimate. The spatial null model was found to be optimum for *Culex* spp. which can be a product of species with high abundances often explaining themselves through spatial variation alone (Gotelli and Ulrich 2012)(Supplementary Table 1). There is a clear pattern of increased abundance along the course of the Galana-Sabaki River which skirts the north boundary of the Arabuko Sokoke National Reserve before entering the sea north of Malindi town. It is visible in the *Culex* model, without covariates, the prediction of the standard error estimates are highest in areas farthest from the sampled point, reflecting the decay of the spatial random effect.

**Figure 5.**
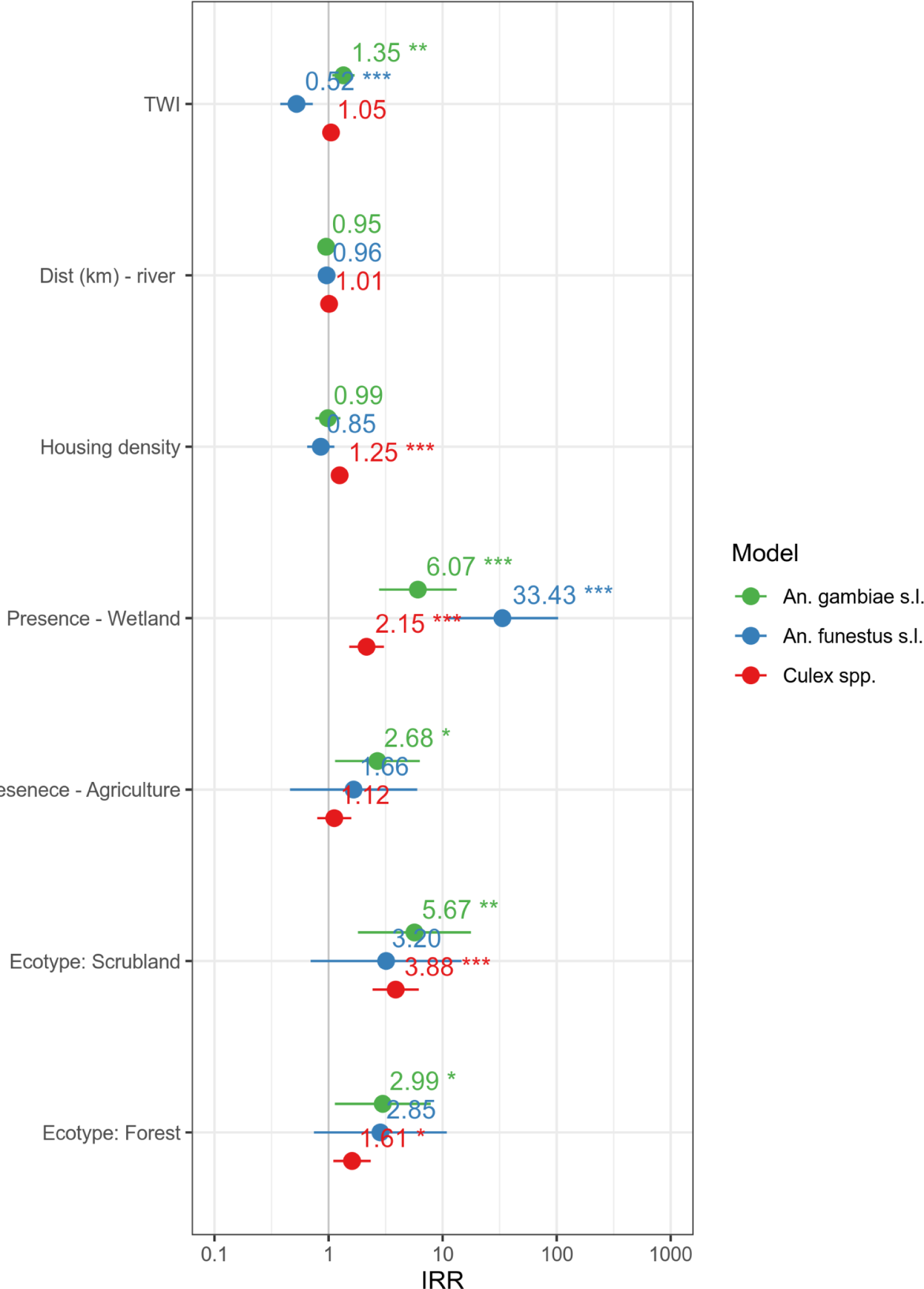
Incidence risk ratio for explanatory variables for household mosquito abundance for the 3 primary species of interest derived from the non-spatial negative binomial GLMM with a random effect for repeated sampling at the household level.

**Figure 6.**
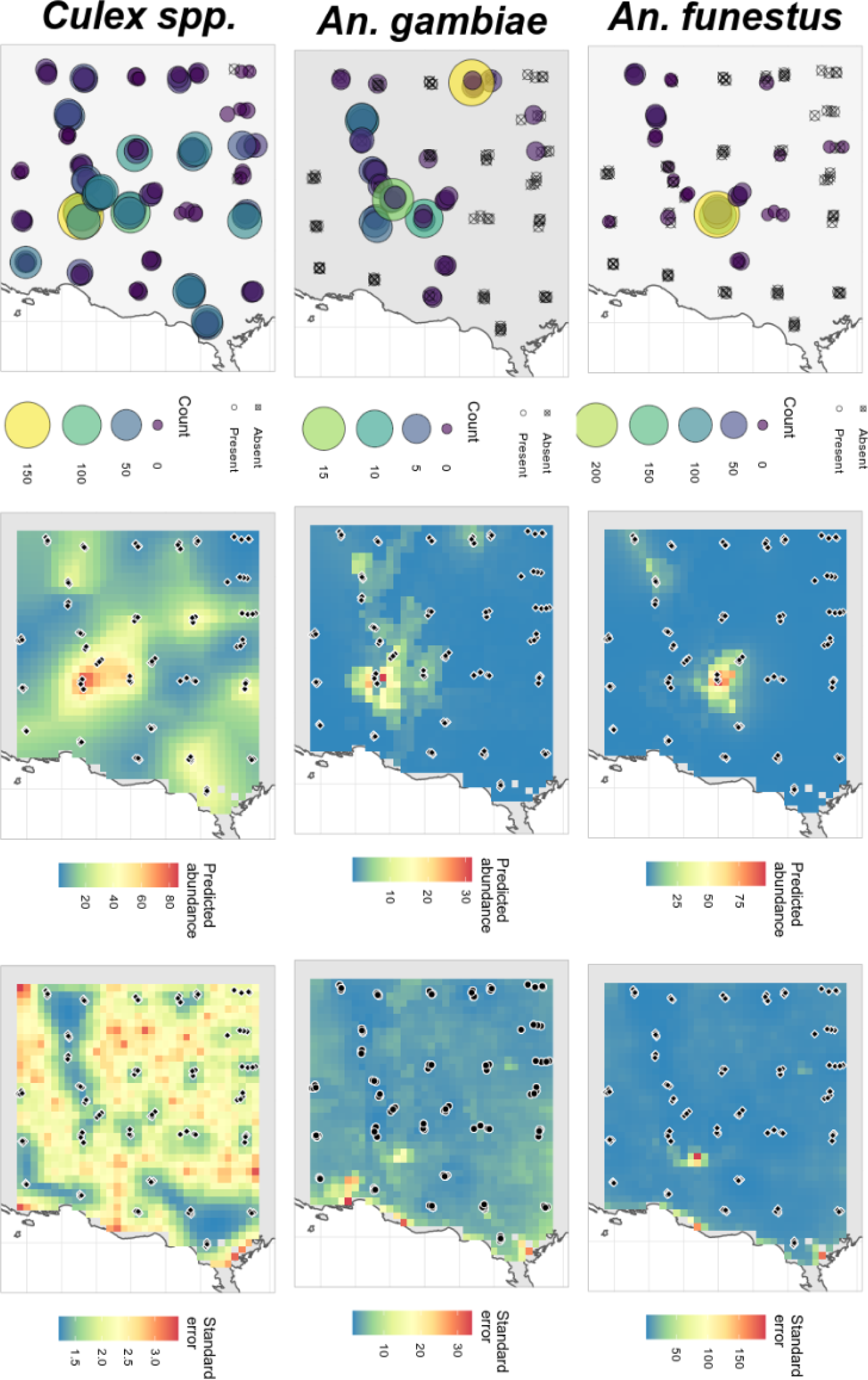
– Observed and modelled species distributions for *An. gambiae* s.l., *An. funestus* and *Culex* spp. The first panel shows the cumulative catch for each house sampled in each sampling location. Black crosses indicate houses that were sampled but none of that species was present. The centre panel is the model estimate created using a combination of the fixed effect and the spatial random field, the associated standard error is shown on the right

## Discussion

In this study we describe the implementation of a reproducible, spatially-explicit sampling approach that can be deployed when surveying a hitherto unstudied region. This relatively short but intensive collection period can aid in identifying regions with high vector (or nuisance biter) abundance that may benefit from targeted interventions such as IRS or larval source management. The rapid characterisation of species diversity, abundance and associated variance facilitates the design of vector control implementation trials with entomological endpoints. The standard error map associated with predictions can also highlight areas in need of further sampling points in future rounds to improve the model’s overall predictive capacity (Kabaghe et al. 2017). Four households were selected as outlined in (Sedda et al. 2019) to reflect the diminishing improvement in reducing the standard error of prediction produced by sampling additional households. The absence of any *Anopheles* spp. mosquitoes in almost half of the sampling points over a relatively small region show the challenges one may expect when designing trials with an entomological endpoint. This can be of particular concern in the use of entomological outcomes in vector control tool trials where defining clusters often occurs without prior entomological data. This can exacerbate heterogeneity in the endpoints making it punitively expensive to power trials or lead to a substantial imbalance between trial arms (Van Hul, Braks, and Van Bortel 2021).

Given the reliance of mosquitoes on water bodies for larval breeding sites, the relationship with wetlands for the two major malaria vector species was not unexpected. However, it does show that satellite-derived data on the presence of wetlands may be incorporated into the exploratory stages of entomological monitoring of the two major malaria species in the area and can provide input that would substantially narrow the area of implementation for a potential larviciding campaign. This strong association with water bodies mirrors previous findings in the region where wind direction and proximity to larval breeding sites were key determinants for malaria transmission in an area (Midega et al. 2012). In addition, abundance collections, while only encompassing part of the risk profile in malaria transmission, can be useful to assess intervention efficacy in pragmatic intervention trials over large regions when a sufficient number of clusters are screened and baseline data is present (Staedke et al. 2020).

The inclusion of *Culex* spp. abundance in this analysis may provide limited direct benefit to an NMCP in this region given the limited concern of disease transmission by *Culex* spp. in this area (Njenga et al. 2017; Grossi-Soyster et al. 2017). However, the assessment of *Culex* spp. presence and density act as useful quality assurance tools for areas where no *Anopheles* were captured that the trap was functioning, and it has also been found that the presence of high Culex spp. densities can have implications on both bednet usage and perception of intervention success. (Brown et al. 2021; Irish et al. 2008).

The targeting of intervention toward regions of high vector abundance has shown some promise with the use of *An. funestus* risk map delineating areas to be included as part of an IRS campaign in Zambia. This targeting of control leads to a greater reduction in malaria incidence than using the traditional administrative unit programme or health facility targeting (Larsen et al. 2020). This study depended on ecologically derived risk maps that are predictive of areas of potential presence, as opposed to areas with high vector abundance, but does illustrate how an improved understanding of mosquito distribution can enhance control efforts.

Our environmental stratification approach coupled with the lattice with close pairs design established a clear relationship with the two primary vector species and the proximity to water and allowed the generation of maps that could aid in refining future sampling or control efforts. Studies on mosquito avoidance behaviour in human populations were conducted in Malindi in the early 2000s where the authors developed a rural-urban stratification to account for the varying factors that can influence mosquito abundance and human-mosquito interaction such as building type, sanitation and socio-economic status (Macintyre et al. 2002; Keating et al. 2003, 2004). These classifications were carried forward by (Mwangangi et al. 2012) and they characterised the major vectors in Malindi town and surrounding environs. They similarly found that *An. gambiae* was localised to the edge of their study area in close proximity to the river. It is uncertain whether run-off from the river of human-mediated activity may be driving this link. Whilst Mwangangi et al. described an urban-rural gradient in their work, their entire research area is encompassed within the urban classification in our sampling strata. This reflects one of the limitations of conducting ecological stratification across such a wide area as the variation encompassed within the resulting ecotype is governed by the scale of the area selected and the spatial resolution of the data used to partition the ecological variation. There is a clear risk that increasing spatial scale in the species abundance modelling models, will increase the potential to both overestimate the low values and underestimate regions of high abundances as a result of insufficient resolution to untangle local regions of high environmental heterogeneity (Waldock et al. 2022).

This work provides a limited snapshot and would need to be expanded to include sampling year-round to account for the temporal variation caused by seasonality. The absence of household characteristics precluded the analysis testing for associations with factors that can be linked to mosquito heterogeneity at a household level (roof type, wall type, presence of open eaves), however, there currently exists limited data on this in a wide spatial scale which would limit the applicability of the approach outside the study area and make predictions impossible in unsurveyed sites (Karisa et al. 2022; Tusting et al. 2017). Additionally, the use of morphological species identification precludes the distinction of *An. gambiae* s.s. and *An. arabiensis* which have well-known bionomic differences. Equally, the *An. funestus* s.l. may be composed of a number of species with varying transmission potential based on their feeding preferences which include *An. rivulorum, An. longipalpis* C, and *An. funestus* s.s. all of which have been detected in outdoor collections in Kilifi county (Kinya et al. 2022). In terms of operational challenges, the requirement to identify four households in close proximity proved challenging in rural areas leading some sites to have a greater distance between households. Due to the poor road network, often the nearest household by straight line distance would not be the easiest accessible household. One solution could be to cross-check the selected points with habitation indices and travel friction datasets prior to deploying field teams to highlight existing road structures and the nearest area of habitation (Longbottom et al. 2020). This would ensure easy identification of homes for sampling and reduce some of the associated costs of carrying out sampling over an expansive area.

## Notes

### Competing Interest Statement

The authors have declared no competing interest.

