## Supplementary Material for "Spatially explicit sampling frameworks to identify regions of increased mosquito abundance"

**Supplementary figures and tables**

**Supplementary Table 1.** Characteristics used to define the three ecotypes classifications in the region including the predominate landcover and the segmentation of the distinguishing climatic variables into low, medium or high values. EVI - Enhanced Vegetation Index; Temp - Temperature; ET - Evapotranspiration.

|  | **Forest** | **Scrubland** | **Urban** |
| --- | --- | --- | --- |
| **Predominate**  **Landcover** | Forest | Mixture of Shrubland, Grassland and Wetland. | Mixture of Water bodies and Urban. |
| **Low** |  | EVI amplitude  Temp variance  Precipitation | ET - mean,  EVI - mean and amplitude  Temp - variance  Precipitation |
| **Medium** | ET - mean, amplitude and variance  EVI - mean, variance and amplitude  Temp - mean and variance  Elevation  Precipitation | ET - mean and amplitude  EVI - mean and variance  Temp - mean and variance | ET - amplitude  EVI - variance |
| **High** | Temp - amplitude | Temp - mean  ET - variance | ET - variance  Temp - mean |

**Supplementary table 2.** Sum predictive log-likelihoods from k-fold cross-validation of the spatial predictive model with the intercept only (null), the variables found to be significant in the non-spatial model (significant only) and all the variables screened (maximal). The lowest negative log-likelihood produces the best fold predictions.

|  | *An. gambiae* | *An. funestus* | *Culex spp.* |
| --- | --- | --- | --- |
| Null | -179.9228 | -176.8462 | -444.0091 |
| Significant only | -159.9427 | -172.3542 | -467.0827 |
| Maximal | -154.4305 | -198.5634 | -467.2399 |

**Supplementary table 3.** Parameter estimates for fixed and random effects from the optimum model for prediction of mosquito abundance. Parameters associated with the GMRF in bold.

| **Parameter** | *An. gambiae* | *An. funestus* | *Culex spp.* |
| --- | --- | --- | --- |
| Intercept | -6.24 (-12.3 - -0.22) | -0.2 (-9.18 - 8.77) | 2.41 (1.8- 3.02) |
| Presence - Wetland | 0.99 ( -0.38 - 2.35) | 1.22 (-0.7 - 3.3) |  |
| TWI | 0.32 (-0.02 0.65) |  |  |
| Presence - - Agriculture | 0.33 ( -1.32 - 1.98) |  |  |
| Dist (km) - river | -0.02 (-0.11 - 0.07) |  |  |
| Housing density | -0.16 (-0.58 0.25) |  |  |
| Ecotype: Scrubland | 0.68 (-1.4 - 2.76) |  |  |
| Ecotype: Forest | 0.19 (-1.7 - 2.08) |  |  |
| **Range (km)** | 13.16 (4.56 - 37.96) | 11.42 (5.29 - 24.6) | 7.36 (3.98- 13.6) |
| **φ** | 2.13 | 1.83 | 3.03 |
| **SD (spatial process)** | 1.44 | 2.3 | 1.2 |
